# DeepDiffusion: a Physics-Informed Neural Network for Heterogeneous Facilitated 1D Diffusion

**DOI:** 10.64898/2026.05.27.727946

**Authors:** Korak Kumar Ray, Benjamin Ambrose, Gabriel della Maggiora, Nicolas de Diego Pinedo, Stefan Bauer, Artur Yakimovich, David S. Rueda

## Abstract

Single-molecule fluorescence microscopy combined with optical tweezers has enabled the direct observation of diffusing proteins on tethered DNA. These and complementary techniques reveal facilitated one-dimensional diffusion as a common functional mechanism for numerous DNA-binding proteins, with a wide range of heterogeneous diffusive behaviours arising from different DNA-binding modes. However, detailed investigations have been limited by the lack of methods to detect such heterogeneous diffusion. We have developed DeepDiffusion, a physics-informed neural network model for estimating the instantaneous diffusion at each point along a single-molecule trajectory. We show, using synthetic trajectories, that DeepDiffusion can accurately detect subtle changes in diffusion even when challenged with large underlying errors. DeepDiffusion can recapitulate previously characterised heterogeneity in experimental data and reveal mechanistic details of facilitated diffusion that are inaccessible to current methods. We expect DeepDiffusion to become a powerful tool for single-molecule researchers, allowing them to investigate the diffusion of their proteins of interest in unprecedented detail.

## Introduction

Facilitated diffusion along one-dimensional (1D) tracks is a ubiquitous feature across a range of biomolecular processes. First implicated for the anomalously fast target binding of the lac inhibitor [1], facilitated 1D diffusion on DNA represents a critical mechanism by which proteins reduce their target search dimensionality within vast cellular genomes [2, 3]. Protein diffusion on DNA arises out of an interplay of specific and non-specific interactions. ‘Sliding’ proteins interact with DNA without dissociating, whilst ‘hopping’ proteins transiently dissociate and reassociate. Thus, facilitated diffusion studies are a powerful tool for interrogating fundamental protein-DNA interactions [3].

Single-molecule microscopy enables the direct imaging of facilitated 1D diffusion [4–7]. Recent advances in correlated optical tweezers with fluorescence microscopy (CTFM) [8] greatly improve our ability to observe protein diffusion on DNA. In CTFM, a labelled protein diffusing on a DNA molecule tethered between optically trapped beads is tracked by fluorescence microscopy (Figure 1a). The resulting kymographs (Figure 1b) reveal the protein’s motion in real-time. Using CTFM, facilitated diffusion has been observed in various contexts [9–19] (Figure 1c).

**Fig. 1.**
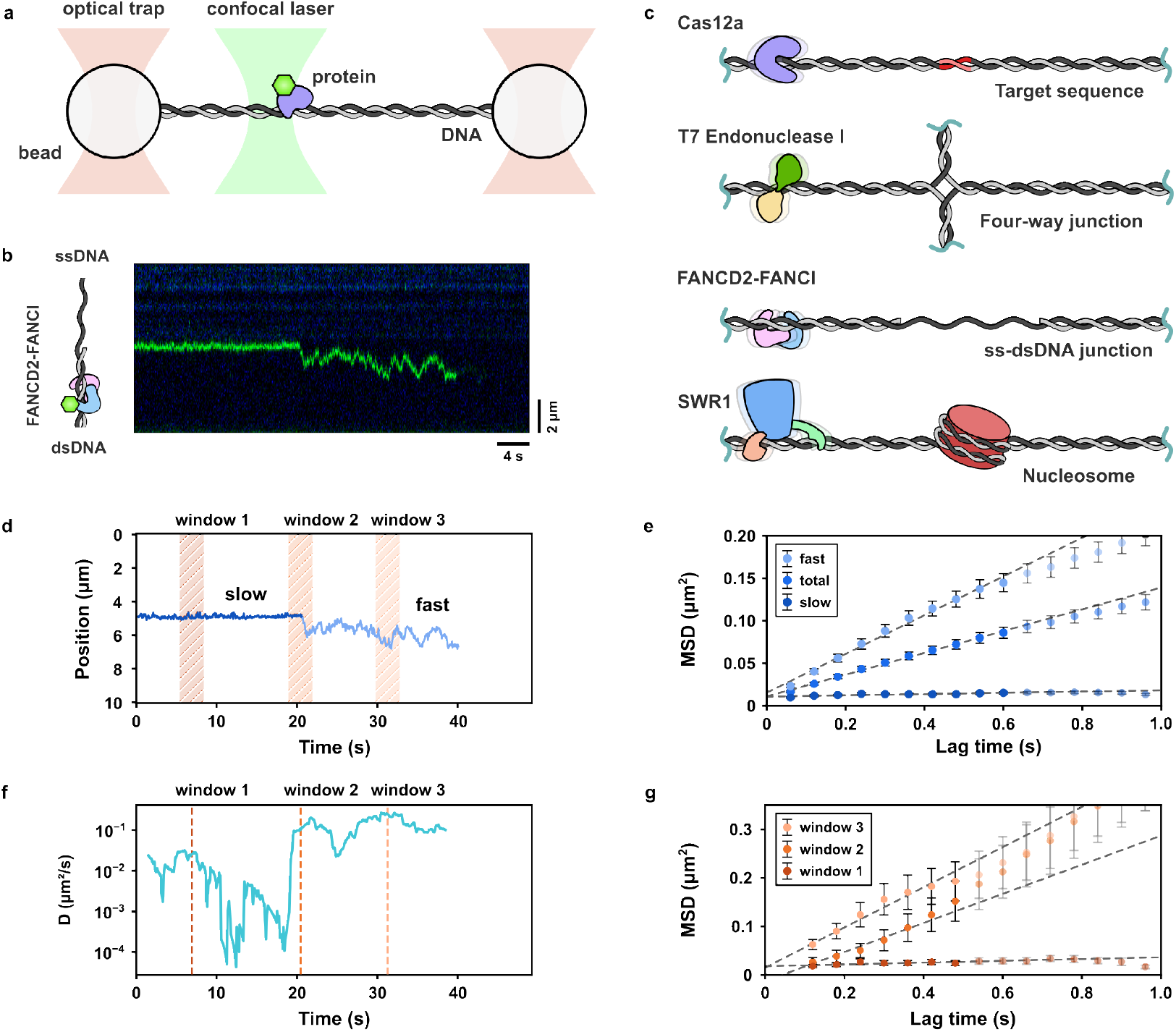
Tracking heterogeneous facilitated 1D diffusion by CTFM. **a**, CTFM schematic with tethered DNA between optically trapped beads scanned with a confocal laser, following a single labelled protein. **b**, Kymograph of labelled FANCD2-I (green) on DNA, transitioning between a bound state and fast diffusion. **c**, Examples of target search for DNA-binding proteins by facilitated 1D diffusion. **d**, Position-time trajectory of FANCD2-I on DNA (from **b**), manually segmented into slow (dark blue) and fast (light blue) phases. Three 50-frame-long windows are shown. **e**, MSD-lag time plots for the fast phase, total trajectory, and slow phase, along with linear fits yielding the diffusion coefficients (mean*±* error of fit) 11.4 *±*0.4 *×*10^*−*2^ *µ*m^2^/s, 6.4 *±*0.1 *×*10^*−*2^ *µ*m^2^/s, 0.4 *±*0.1 *×*10^*−*2^ *µ*m^2^/s, respectively. Error bars correspond to standard errors of mean (s.e.m). **f**, Instantaneous *D* trajectory from **d** using the rolling-window method. Dashed lines depict windows in **d. g**, MSD-lag time plots for the three windows in **d**, along with linear fits. Error bars correspond to s.e.m.

The protein’s fluorescent signal is used to calculate a single-molecule time trajectory (Figure 1d). The mean-squared displacement (MSD, Figure 1e) is linearly fit to determine a single diffusion coefficient (*D*) for the trajectory [20]. However, CTFM and other single-molecule experiments have revealed heterogeneous diffusion that cannot be explained by a single *D* value [12, 18, 21, 22]. Detecting and measuring changes in diffusion, though, has proven challenging. Non-interchangeably heterogeneous trajectories (Figure 1e) can be manually segmented to determine different *D*s [10, 16]. A rolling-diffusion approach can further detect multiple interchangeable *D*s in a single trajectory without manual segmentation [12, 18] (Figure 1f and g). However, this is limited by the *ad hoc* rolling window size: small windows lead to less precise *D*s, whereas larger windows blurs transitions. Furthermore, this need to optimise window sizes makes comparisons across datasets and experiments challenging. Thus, quantifying heterogeneous facilitated diffusion is limited by conventional analysis methods.

Recent advances in deep-learning-based (DL-based) approaches to signal processing of diffusion trajectories [23, 24] and inverse problems in microscopy [25–27] show that a highly expressive data-driven deep neural network (DNN) can facilitate accurate solutions to analysing heterogeneous facilated 1D diffusion. Furthermore, these solutions can be improved by incorporating the underlying physics [28, 29]. However, existing approaches [30] are typically optimised for two-dimensional (2D) microscopy data [31] or focus on classifying non-Brownian diffusion types [32, 33].

Here, we have developed DeepDiffusion, a physics-informed neural-network-based analysis pipeline to estimate instantaneous diffusion coefficients from single-molecule trajectories. Using synthetic trajectories, we find that DeepDiffusion yields significantly more precise and accurate estimates of heterogeneous instantaneous diffusion coefficients than current approaches. Applying DeepDiffusion to experimental data, we show that it outperforms rolling-window analysis in detecting changes in instantaneous diffusion coefficients. We have packaged DeepDiffusion into an easily executable, open-source analysis pipeline. We anticipate DeepDiffusion to be an important asset for single-molecule analysis of facilitated 1D diffusion, enabling quantification of heterogeneous diffusion with greater precision and accuracy.

## Results

### Architecture Design for DeepDiffusion

The main challenge in quantifying *D* from single-molecule trajectories stems from localisation errors (*LE*s) that obscure the true protein location. To design a model architecture that determines instantaneous *D* from single-molecule trajectories, we considered several experimental noise sources. We use a single *LE* term that captures (i) the apparent protein motion arising from bead diffusion, (ii) the force-dependent DNA tether fluctuations, and (iii) limited localisation precision due to low photon counts. We model *LE* as a Gaussian distribution centered at the ‘true’ protein location.

We designed DeepDiffusion to distinguish between displacement caused by free diffusion and apparent displacement due to *LE*. Unlike the uncorrelated free diffusion displacements, apparent displacements caused by *LE* are correlated [34]. Feeding position-time trajectories (e.g., Figure 1d) into a series of 1D convolution blocks with subsequent long short-term memory (LSTM) layers (Figure 2a) allows us to differentiate between *LE* and diffusion, yielding the diffusion (*g*_*ϕ*_(*t*)), drift (*f*_*θ*_(*t*)) and learnable reweighting (*U*_*v*_(*t*)) terms (Figure 2b).

**Fig. 2.**
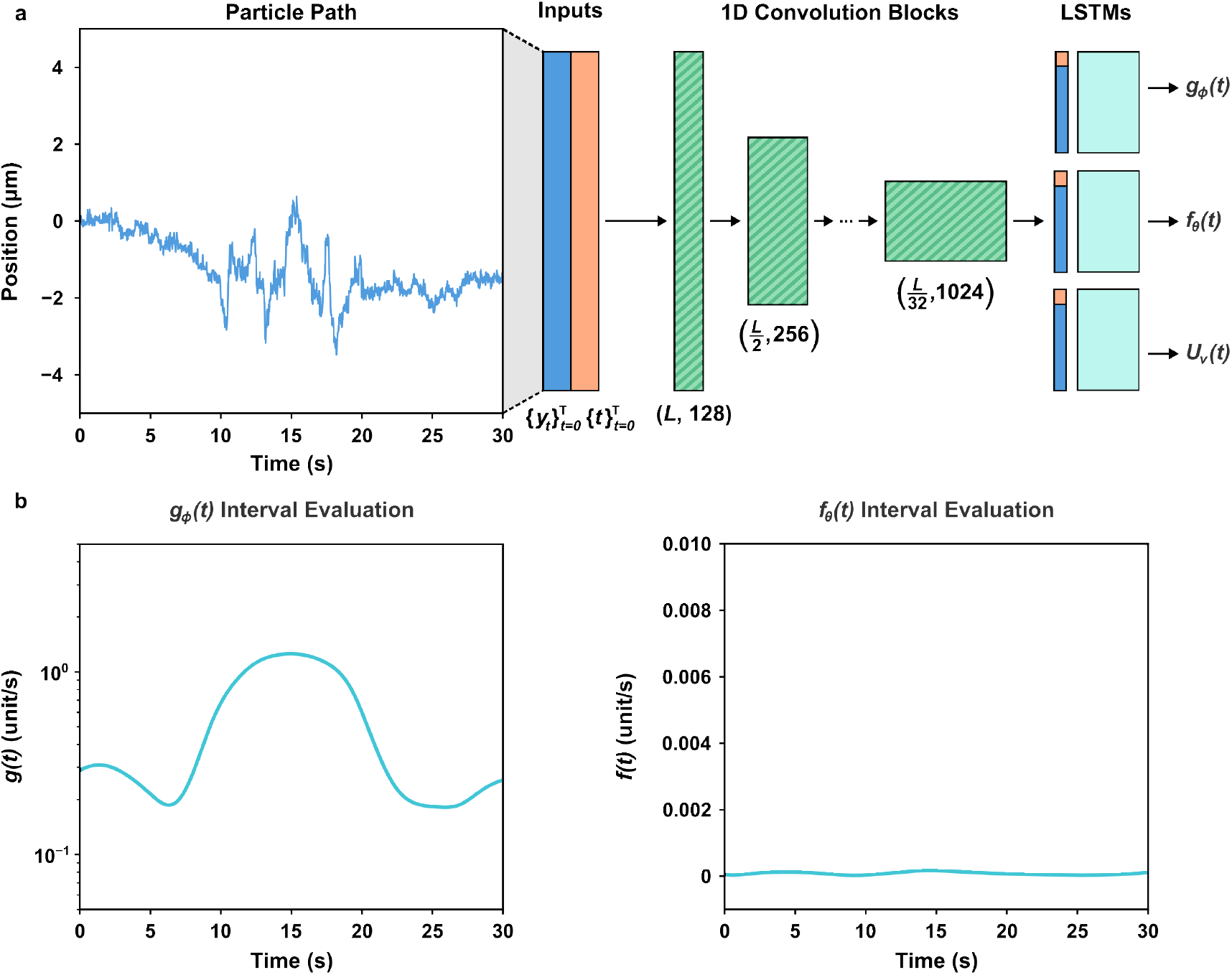
Model Architecture. **a**, The conditional SDE inference architecture schematic, where the observed noisy trajectory and its time grid are encoded with a 1D residual CNN with temporal pooling, then summarized by a unidirectional LSTM into a path embedding [*X*] that conditions the time-dependent diffusion (*g*_*ϕ*_([*X*], *t*)), drift (*f*_*θ*_ ([*X*], *t*)), and the learnable reweighting (*U*_*v*_ (*t*, [*X*])) terms. **b**, Interval evaluations of the model outputs for the same input, showing the predicted diffusion *g*_*ϕ*_([*X*], *t*) (*left* ) and drift *f*_*θ*_ ([*X*], *t*) (*right* ) terms.

### Evaluating DeepDiffusion using synthetic homogeneous datasets

To test the ability of DeepDiffusion to estimate instantaneous *D*, we generated synthetic datasets of single-molecule trajectories with single constant *D* and *LE* in typical experimental ranges (respectively, 10^*−*4^–1 µm^2^/s and 0 – 300 nm, Figure 3a). DeepDiffusion can robustly predict the “true” instantaneous *D* for these trajectories even in the presence of *LE* (Figure 3a). With increasing *LE*, the predicted *D* distributions widen (Figure 3b). This widening is asymmetric, and is biased towards lower *D* predictions (Figures 3c and 4a). A likely explanation for this error is our model’s inability to distinguish between “true” displacements and apparent displacements arising from *LE* in the low-*D* and high-*LE* regimes (see Discussion). However, for typical experimental *LE* ranges (*≤*100 nm), DeepDiffusion predicts accurate instantaneous *D*s over more than three orders of magnitude, encompassing the most commonly encountered experimental range.

**Fig. 3.**
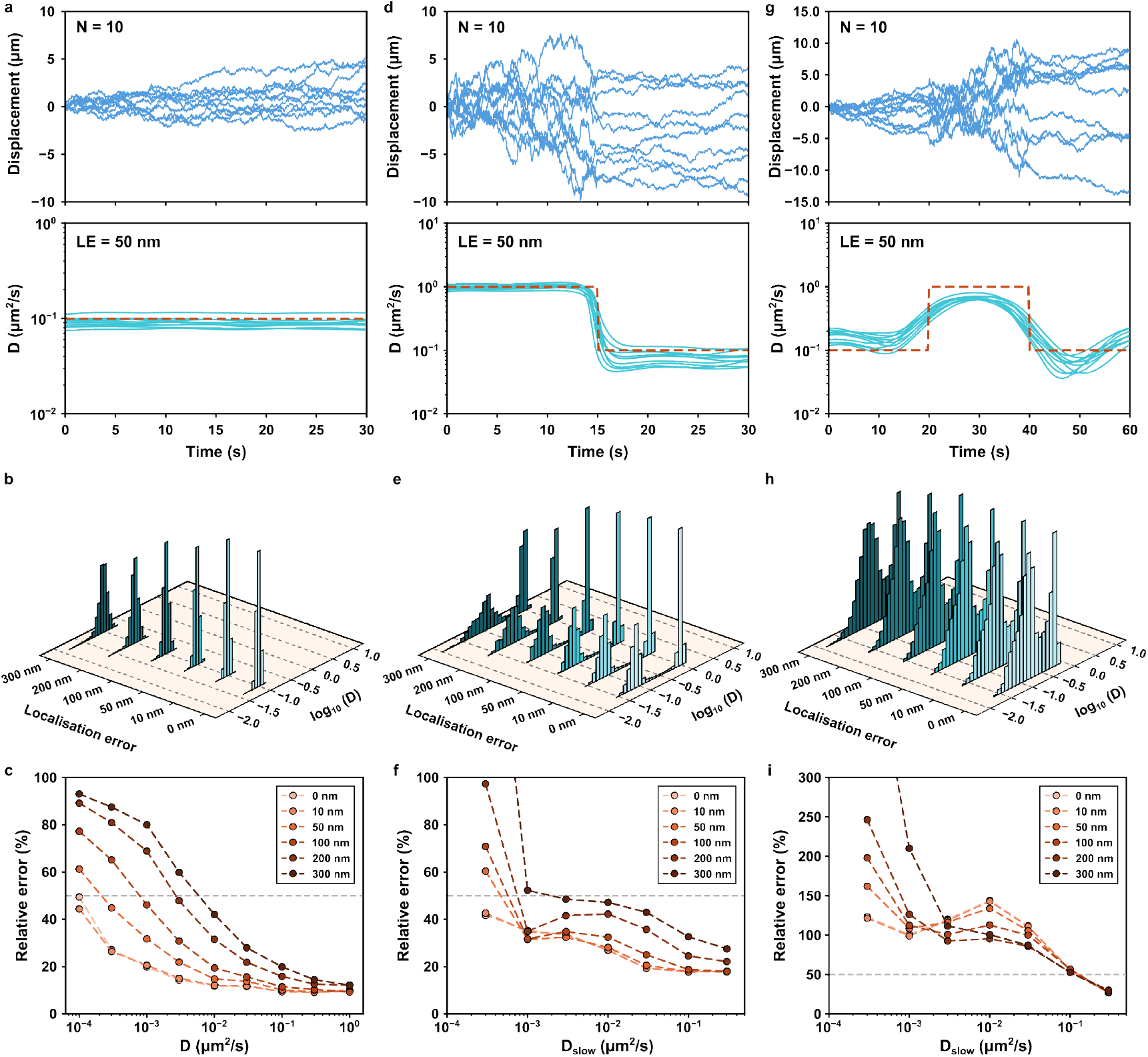
Evaluation of DeepDiffusion using synthetic datasets. **a**, Representative single-molecule trajectories (blue) with constant *D* (*top*), with predicted (cyan) and true (red) instantaneous *D* (*bottom*). **b**, Predicted *D* histograms with increasing *LE* for *D* = 0.1 µm^2^/s. **c**, Relative errors of predicted *D* plotted for varying *D*s and *LE*. **d**, Representative single-molecule trajectories (blue) with a single change in *D* (*top*), along with predicted (cyan) and true (red) instantaneous *D* (*bottom*). **e**, Predicted *D* histograms with increasing *LE* for initial *D*_*fast*_ = 1 µm^2^/s and final *D*_*slow*_ = 0.1 µm^2^/s. **f**, Relative errors of predicted *D* plotted for varying final *D*_*slow*_s and *LE*. **g**, Representative single-molecule trajectories (blue) with two changes in *D* (*top*), along with predicted (cyan) and true (red) instantaneous *D* (*bottom*). **h**, Predicted *D* histograms with increasing *LE* for an initial and final *D*_*slow*_ = 0.1 µm^2^/s and an intermediate *D*_*fast*_ = 1 µm^2^/s. **i**, Relative errors of predicted *D* plotted for varying intermediate *D*_*slow*_s and *LE*.

Next, we compared DeepDiffusion to the rolling-window approach for the same homogeneous datasets. For *LE* = 0, the rolling window method and DeepDiffusion yield similar results (Figure 3c and Supplementary Figure S1c). However, the rolling window precision drastically worsens with nonzero *LE*, especially at lower *D* values. For high *LE*, a spurious peak appears in the instantaneous *D* distribution greater than the ground truth (Supplementary Figure S1b), resulting in plateauing mean estimated *D*s (Figure 4a). This arises from the rolling window method fitting the noise from *LE* rather than the diffusion. Even when the rolling window method correctly estimates the mean *D*, we observe a significant number of large (over 10-fold) false-positive *D* transitions (Supplementary Figure S1a and b). By comparison, DeepDiffusion yields more stable predictions of instantaneous *D*, thereby reducing the chance of such spurious transitions.

**Fig. 4.**
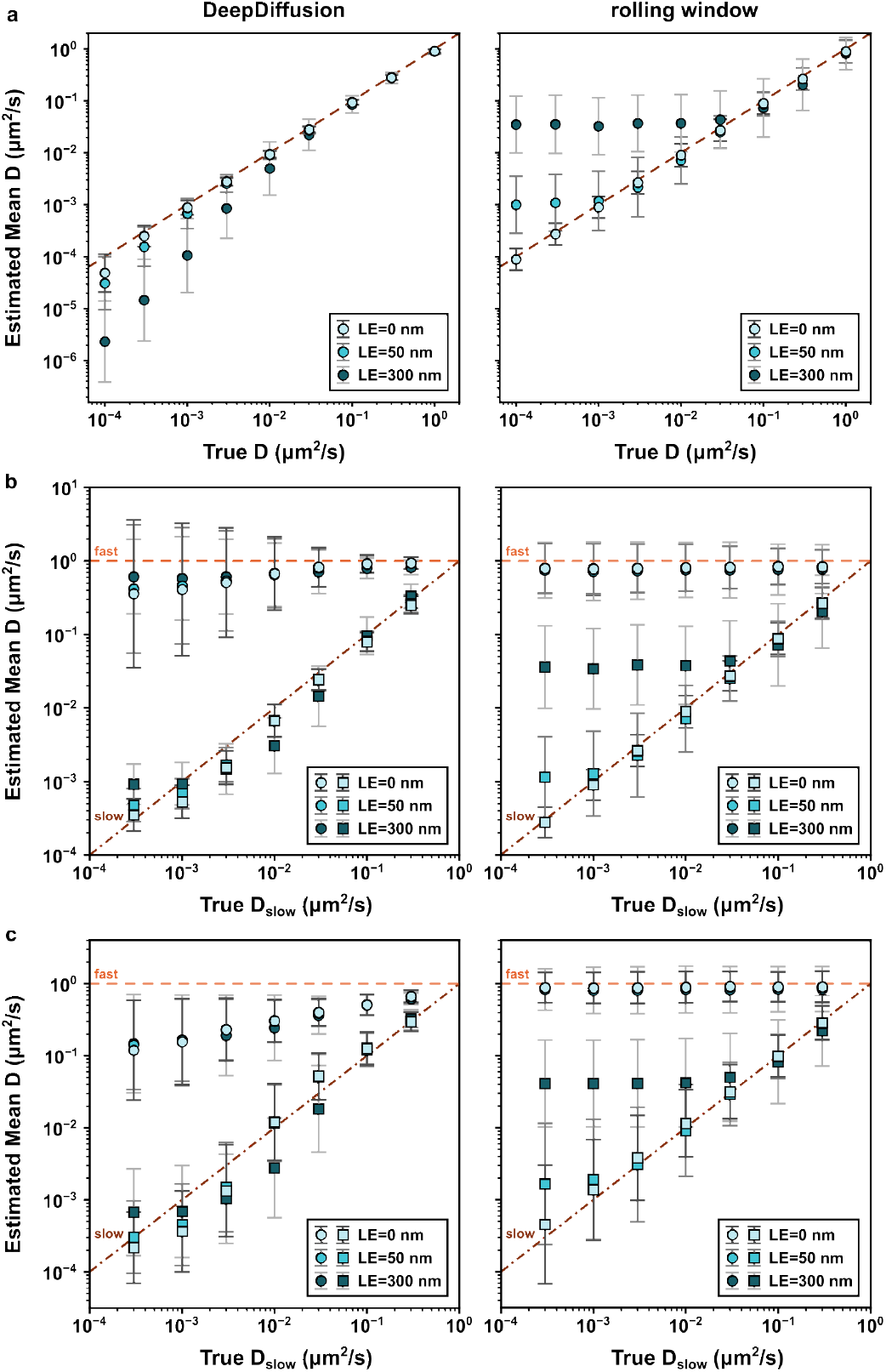
Evaluation of DeepDiffusion bias using synthetic datasets. **a**, Mean *D* calculated using DeepDiffusion (left) and rolling window (right) for constant *D*, plotted as a function of the true *D* for different *LE*s. The dashed line *y* = *x* represents the case of no overall bias. **b**, Mean *D*s (circles and squares for *D*_*fast*_ and *D*_*slow*_, respectively) calculated using DeepDiffusion (left) and rolling window (right) for a single-step decrease from a constant higher *D*_*fast*_ to a varying lower *D*_*slow*_, plotted as a function of the true *D*_*slow*_ for different *LE*s. The dashed lines *y* = 1 and *y* = *x* represent the cases of no overall bias for *D*_*fast*_ and *D*_*slow*_ respectively **c**, Mean *D*s (circles and squares for *D*_*fast*_ and *D*_*slow*_, respectively) calculated using DeepDiffusion (left) and rolling window (right) for a two-step change from a varying lower *D*_*slow*_ to a constant higher *D*_*fast*_ and back, plotted as a function of the true *D*_*slow*_ for different *LE*s. The dashed lines *y* = 1 and *y* = *x* represent the cases of no overall bias for *D*_*fast*_ and *D*_*slow*_ respectively. Error bars correspond to standard deviations of the *D* distributions.

### Evaluating DeepDiffusion using synthetic heterogeneous datasets

We next investigated the performance of DeepDiffusion on synthetic datasets exhibiting changes in *D*. First, we generated datasets where the instantaneous *D* showed a single-step decrease from a constant higher *D*_*fast*_ to a varying lower *D*_*slow*_ (Figure 3d). DeepDiffusion can distinguish these two diffusive states over mutliple folds changes in *D* for a large range of *LE* (Figure 3d and e). While the overall precision is lower for this heterogeneous dataset than the above homogeneous one, Deep-Diffusion still returns more precise instantaneous *D* estimates than the rolling window in all cases (Figure 3f and Supplementary Figure S2c). Similar to the homogenous dataset above, increasing *LE* leads to increasing lower *D* predictions for DeepDiffusion (Figure 4b). However, compared to the rolling window method (Supplementary Figure S2a and b), DeepDiffusion is less likely to predict spurious instantaneous *D* transitions for this dataset.

Unlike the homogeneous case, the spurious lower *D* predictions at high *LE* does not lead to an underestimation of the mean *D*s for the two states (Figure 4b). Instead, DeepDiffusion yields a slight bias in estimating the mean *D*s independent of *LE* for a large (greater than two orders of margnitude) change in *D*. This estimation bias leads to the higher error at these larger changes in *D* (Figure 3f). In contrast, the rolling window method shows an *LE*-dependent plateauing at low values of *D*_*slow*_, similar to the homogeneous case.

Surprisingly, we observe a direction dependence in DeepDiffusion. Using DeepDiffusion on a heterogeneous dataset where the instantaneous *D* shows a single-step increase from a varying lower *D*_*slow*_ to a constant higher *D*_*fast*_, we see a different response to the single-step decrease dataset (Figure 3f and Supplementary Figure S3c). In contrast, the results for the two datasets using the rolling window method are identical (Supplementary Figure S2). This demonstrates that the difference does not exist in the test datasets, and arises due to DeepDiffusion instead. Regardless of this slight difference, however, DeepDiffusion yields more robust and precise instantaneous *D* predictions than the rolling window method for both datasets, especially at moderate to high *LE*.

Finally, we tested DeepDiffusion with more complex datasets showing two transitions in *D* (Figure 3g and Supplementary Figure S3d). In each case, DeepDiffusion can distinguish between the two diffusive states, even in the presence of high *LE*, albeit with a lower precision than the single-transition datasets (Figure 3 and Supplementary Figure S3). This is due to an estimation bias of the mean *D* for the two states (in particular, the state with the higher *Dfast* for large *D* changes). However, even for these larger changes, DeepDiffusion shows a lower dependence on *LE*, unlike the rolling window method, which shows a similar plateauing behaviour as above (Figure 4c and Supplementary Figure S4b).

In all of the tested cases, DeepDiffusion shows a significantly better performance for very small changes in *D* (from a three-fold to a 10-fold change) compared to the rolling window method, which does not yield robust results in this regime, making it a critical area for improvement.

### DeepDiffusion recapitulates D2-I binding in high *LE* imaging conditions

We next tested DeepDiffusion on experimental datasets. The sliding clamp FANCD2-FANCI (D2-I) has been previously shown to diffuse freely on double-stranded DNA (dsDNA) with an average *D ≈* 0.1 µm^2^/s [12]. D2-I binds specifically to single-stranded-DNA-double-stranded-DNA junctions (ss-dsDNA junctions), becoming motionless in the DNA frame-of-reference with all the observed motion arising due to *LE*. D2-I thus serves as a well-characterised model system for two drastically different diffusive behaviours (static and diffusing) (Figure 1). Previous characterisation of the heterogeneous diffusion of D2-I on DNA was performed in low *LE* imaging conditions [12], under forces (∼15 pN) that restrict DNA track motion and with high excitation powers (producing high, 80-100, photon counts per subpixel localization) that yield more precise localisations [35].

We used DeepDiffusion to characterise D2-I diffusion under high *LE* conditions (∼5 pN forces and 10-40 photon counts per subpixel localisation). We specifically chose two trajectories that high-tlight the two different D2-I diffusive behaviours (Figure 5a and b). Based on our results with the synthetic datasets (Figures 3d and 3a), we used four iterations of DeepDiffusion to generate a space- and time-symmetrised instantaneous *D* prediction with corresponding model uncertainties (Figure 5a and b) (see Materials and Methods).

**Fig. 5.**
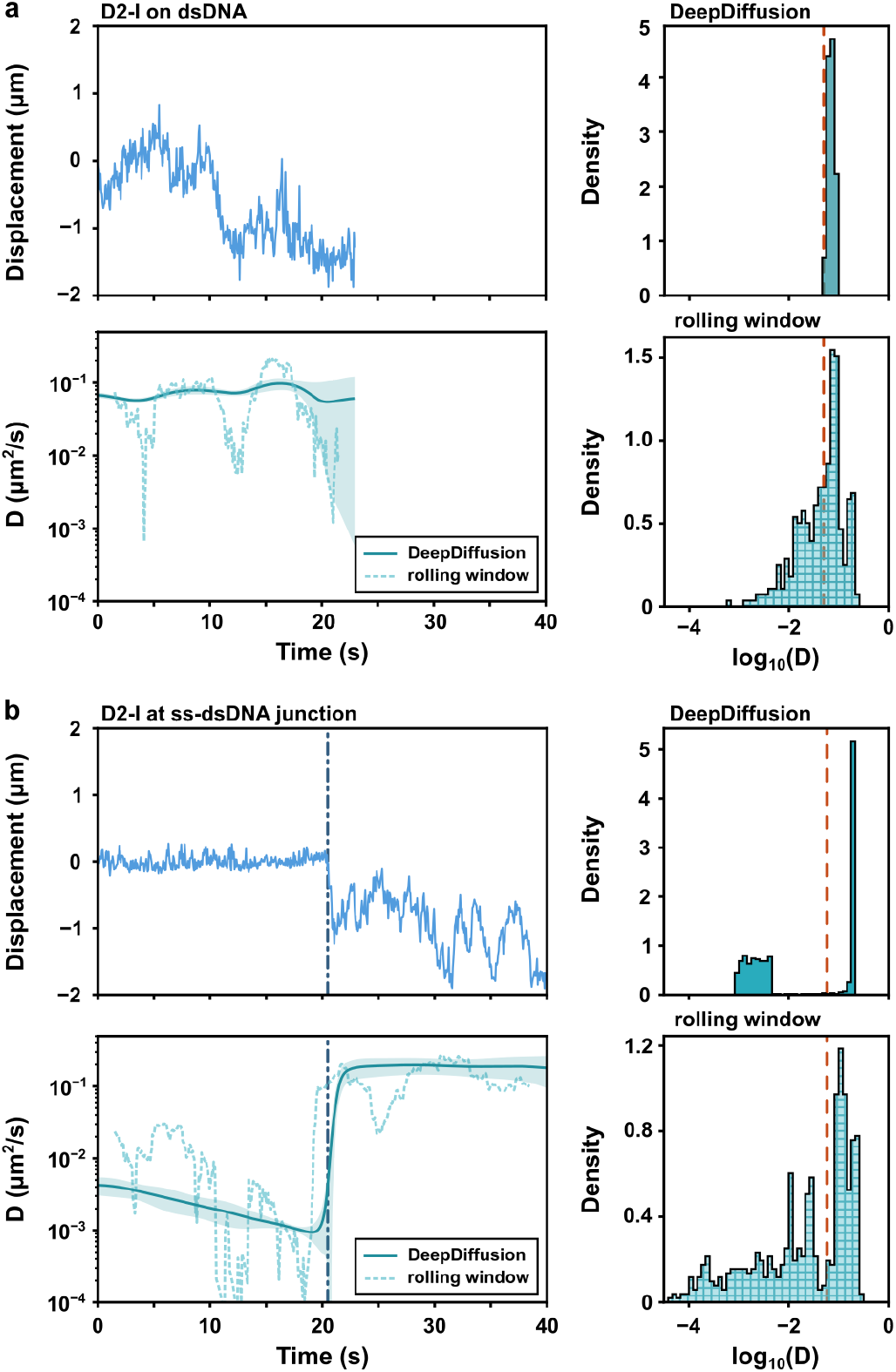
DeepDiffusion distinguishes between bound and diffusing D2-I. (*left* ) Position-time trajectory (*top*) of a D2-I molecule **a**, diffusing on dsDNA, and **b**, bound to an ss-dsDNA junction and diffusing on dsDNA, with the corresponding *D* trajectories (*bottom*) calculated using DeepDiffusion (solid line) and rolling window (dashed line). The shaded region corresponds to the uncertainty in DeepDiffusion. The manually segmented transition between bound and diffusing states in **b** is indicated with the dashed grey line. (*right* ) *D* histograms computed using DeepDiffusion (*top*) and rolling window (*bottom*). The average *D* (corresponding to the MSD *vs* lag-time fit for the whole trajectory) is shown by the dashed red line.

For D2-I diffusing on dsDNA, DeepDiffusion returns a precise *D* trajectory with low model uncertainty (Figure 5a). In contrast, the rolling window method exhibits transient sharp *D* transitions, likely from fitting artifacts due to high *LE*, similar to above (Figure 1). However, a large increase in model uncertainty for the DeepDiffusion predicted *D* is observed towards the trajectory end. This correlates with a similar decrease in the result for the rolling window method. This suggests that there is some evidence in this section of the trajectory for a *D* change. This evidence is picked up by some iterations of the model, but is insufficient for others, likely due to its short-lived nature. In the absence of sufficient evidence, the symmetrised DeepDiffusion output does not yield a change in *D*. However, it returns a large uncertainty in the output, denoting this insufficient evidence.

For D2-I, both static at an ss-dsDNA junction and diffusing on dsDNA, DeepDiffusion robustly differentiates between the static (low *D*) and diffusing (high *D*) states with low uncertainty (Figure 5b). The transition point predicted by DeepDiffusion is identical to the one estimated by manual segmentation (Figure 1d). In comparison, the instantaneous *D* trajectory using the rolling-window method is significantly less precise, with spurious transitions in the bound D2-I state due to low *D* and high *LE* (as seen in the test cases above). The rolling window method further misidentifies the transition point, likely due to averaging between the two diffusive behaviours by windows crossing the transition. By obviating the need for windows, DeepDiffusion thus yields more precise estimations of changing diffusive behaviours (see Discussion).

### DeepDiffusion reveals complex heterogeneous INO80 diffusion

We finally investigated the new mechanistic insights enabled by DeepDiffusion. Facilitated diffusion on DNA, as mentioned above (see Introduction), have been used as a powerful method for investigating protein-DNA interactions. Two major theoretical mechanisms have been proposed for facilitated diffusion, sliding and hopping [3]. Experimentally distinguishing between these two modes is non-trivial and typically achieved by measuring diffusion under varying salt concentrations. A sliding protein maintains constant contact with DNA and thus, the number of protein-DNA interactions remains unchanged during motion. In contrast, a hopping protein microscopically dissociates and re-associates on DNA, thereby breaking and reforming interactions. Therefore, salt-dependent electrostatic screening is expected to affect hopping, but not sliding. Based on this, facilitated diffusion studies characterise proteins with salt-dependent *D* as hopping on DNA [36], while those with salt-independent *D*s are considered to slide on DNA [6].

We tested this model for the chromatin remodeller, INO80 [37–39]. Previous studies on similar chromatin remodellers have shown that these complexes diffuse on dsDNA [17, 18]. In particular, a closely related remodeller, SWR1, diffuses faster on dsDNA (*i*.*e*., with a higher *D*) with increasing salt concentrations, indicating that SWR1 hops on DNA [17].

Our data show that, similar to other chromatin remodellers, INO80 diffuses on dsDNA (Figure 6a and b). The mean trajectory-averaged *D* (*D*_*avg*_) for individual INO80 molecules (calculated using the MSD vs lag-time fit, similar to Figure 1E) under low salt (25 mM KCl) is 0.42 *±* 0.04 µm^2^/s (s.e.m.). Unlike SWR1, the *D*_*avg*_ under high salt (200 mM KCl) remains virtually unchanged (mean *D*_*avg*_ = 0.44 *±* 0.02 µm^2^/s, Figure 6C). This is also reflected in the overall *D*_*avg*_ distributions for these conditions (Figure 6c). Based on the above, this would suggest that INO80 slides on DNA.

**Fig. 6.**
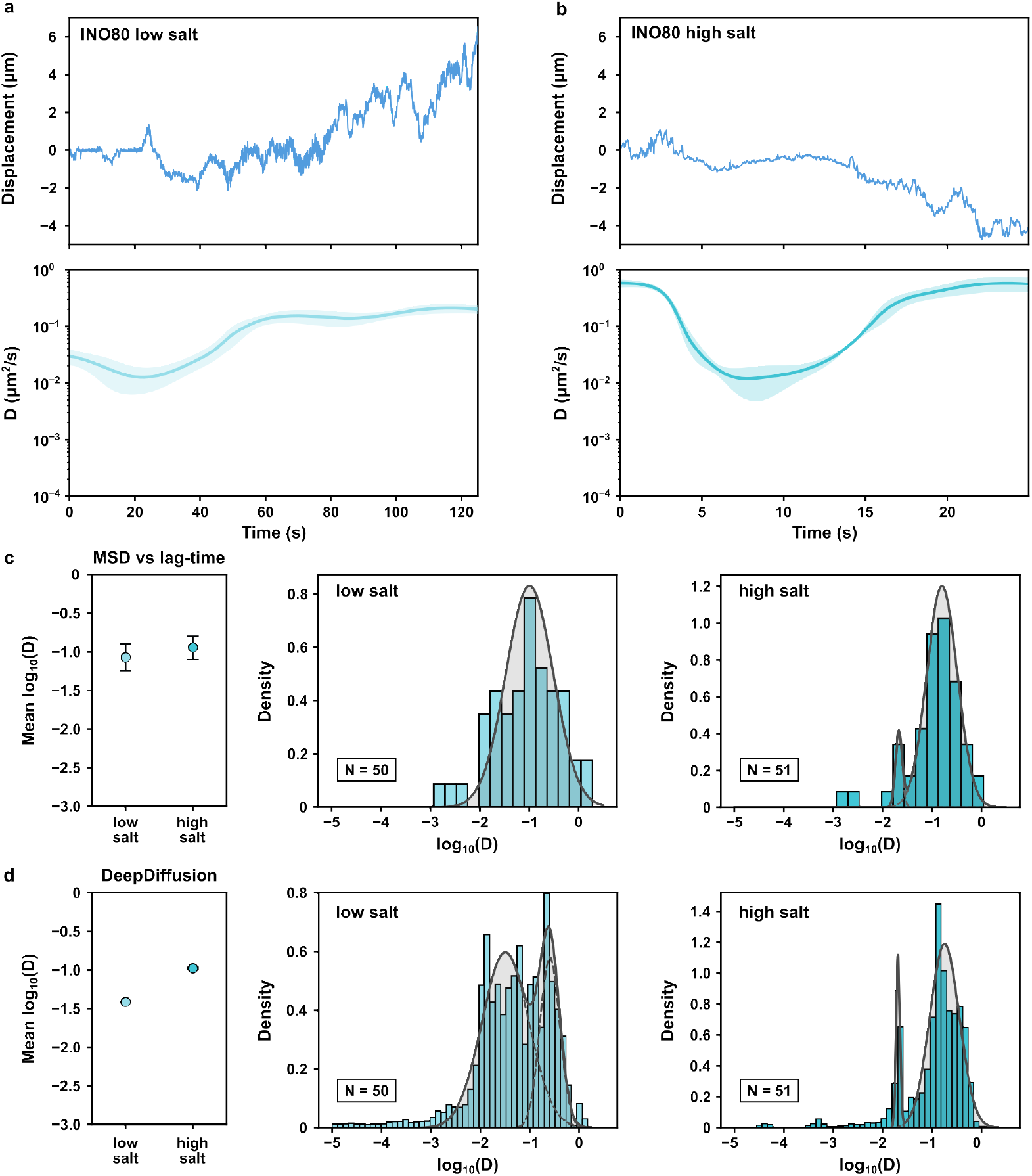
DeepDiffusion reveals salt-dependent changes in diffusion of INO80. (*left* ) Displacement *vs* time trajectory (*top*) of an INO80 molecule diffusing on dsDNA **a**, in low salt, and **b**, in high salt, with the corresponding *D* vs time trajectories (*bottom*) calculated using DeepDiffusion. The shaded region corresponds to the uncertainty in the DeepDiffusion trajectory. **c**, The mean trajectory-averaged *D* (*left* ) for INO80 in low and high salt computed using MSD *vs* lag-time fit for the whole trajectory, and histograms of trajectory-averaged *D* for low salt (*middle*) and high salt (*right* ) conditions. **d**, The mean instantaneous *D* (*left* ) for INO80 in low and high salt computed DeepDiffusion, and histograms of instantaneous *D* for low salt (*middle*) and high salt (*right* ) conditions. Error bars correspond to s.e.m.

We next calculated the instantaneous *D* (*D*_*inst*_) for INO80 using DeepDiffusion. In contrast to the trajectory-averaged results, we observed a salt-dependent increase in the mean *D*_*inst*_ with increasing salt, from 0.327 *±* 0.001 µm^2^/s to 0.435 *±* 0.001 µm^2^/s (Figure 6d), suggesting that INO80 hops on DNA. However, the individual *D* trajectories reveal that INO80 transitions between at least two different diffusive behaviours, a fast diffusing state (*D* ∼ 0.1 µm^2^/s) and a slow diffusing state (*D* (∼ 0.1 µm^2^/s). Both states are present under different salt conditions, and the mean *D* for each state is salt-independent. Instead, it is a change in the relative proportion of each state that drives the salt-dependent increase in the mean *D*_*inst*_ (Figure 6d).

These results demonstrate that INO80 diffusion on dsDNA cannot be characterised by the simple dichotomy of sliding *vs* hopping. Instead, they reveal a far more complex and richer diffusive behaviour of facilitated diffusion on DNA (see Discussion). These insights are only made possible by the robust and precise instantaneous *D* trajectories yielded by DeepDiffusion.

## Discussion

CTFM is a powerful method to directly observe the facilitated diffusion of proteins on DNA. However, CTFM diffusion data analysis is complicated by *LE*. As a comparison, for a frame-time of 30 ms, a particle with a *D* of 0.1 µm^2^/s exhibits a true displacement of ∼60 nm, similar to the apparent displacement of a static molecule due to a *LE* = 50 nm and nearly an order of magnitude lower than the apparent displacement of ∼480 nm for *LE* = 300 nm. Any method for the analysis of instantaneous diffusion must therefore accurately distinguish between the apparent and the ‘true’ displacements of the diffusing molecule.

We have developed DeepDiffusion, a neural network-based analysis pipeline trained to make this distinction. DeepDiffusion can accurately predict instantaneous *D* over a large range of *D* and *LE* (10^*−*4^–1 µm^2^/s and 0–300 nm, respectively). Compared with the rolling window method, DeepDiffusion yields more precise and accurate results on both synthetic and experimental data. In contrast to other methods, DeepDiffusion is most accurate for small *D* changes, with minimal spurious transitions. This is particularly important because these small changes in *D* are expected to encode crucial mechanistic details about protein-DNA interactions that are currently inaccessible. DeepDiffusion detects changes in *D* only for significant evidence of such changes. However, using multiple Deep-Diffusion iterations, we can also capture regions of low transitions evidence through high uncertainty, allowing researchers to assign confidence to the predicted instantaneous *D* trajectories.

Most importantly, DeepDiffusion removes the need for *ad hoc* analysis optimisation. The rolling window method fails to capture true diffusion below a certain threshold and instead captures the noise due to *LE* (Figure 4). Larger window sizes can better differentiate between diffusion and *LE*, but at the cost of lower temporal resolution due to the larger window. Therefore, window sizes need to be optimised for every analysis in an *ad hoc* manner. This further complicates comparing such analyses across different datasets and users. By removing the need for any such optimisation, DeepDiffusion simplifies both the analysis process and the comparisons between diferrent analyses.

Our analysis of heterogeneous D2-I diffusion under high *LE* imaging conditions (i.e., low force and excitation power) highlights the impact of DeepDiffusion on experimental design. CTFM experiments for facilitated diffusion must balance optimised imaging with biologically relevant conditions. For instance, higher forces restrict intrinsic DNA track motions, reducing *LE*, but can affect biomolecular function [40] and nucleic acid structures (such as nucleosomes [41] and junctions [10]). Similarly, increasing photon counts per trajectories increase sub-pixel localisation precision but limit experimental information. For example, higher laser powers decrease trajectory lengths by increasing the photobleaching rate, while downsampling trajectories averages over faster kinetics. Thus, by yielding accurate results under high *LE* imaging conditions (Figure 5), DeepDiffusion enables previously inaccessible facilitated diffusion experiments.

As an example of one such experiment, we used DeepDiffusion to demonstrate how INO80 facilitated diffusion does not fall into the dichotomy of sliding *vs* hopping. INO80 has mutliple DNA binding domains [39], which can lead to different DNA binding modes. One possible rationalisation of our results is that two such binding modes lead to the two observed diffusive states. The constant mean *D* for each state with varying salt suggests that INO80 slides on DNA in both modes (Figure 6). Furthermore, the salt-dependent proportions of the two states indicates that the transition between the two modes depends on the screening of charged protein-DNA interactions. One possible explanation is that dissociation of specific DNA-binding INO80 domains leads to the two different binding modes and diffusive behaviours. A deeper enquiry of the specific structural and mechanistic origins of this heterogeneous diffusive behaviour is, of course, beyond the scope of this study.

We expect such complex diffusive behaviour to be a more general feature of DNA-binding proteins, rather than being limited to INO80 alone. However, such behaviour cannot be easily detected and characterised using trace-averaged *D*s. As we show above, accurate instantaneous *D* trajectories, yielded by methods like DeepDiffusion, are required detect and such quantify heterogenous diffusion. In addition, DeepDiffusion can also identify subtle DNA sequence- and structure-dependent diffusion changes that may be missed by current methods. To enable such in-depth characterisation of facilitated diffusion, we have packaged DeepDiffusion and other tracking analysis tools into a web-based portal called KymoSketch, which we anticipate will be an asset to the single-molecule field.

## Acknowledgments

The authors thank M. Skehan, P. Girvan, and D. Wigley (Imperial College London) for labelled INO80, as well as P. Alcón and L. A. Passmore (MRC LMB) for the use of previously collected D2-I single-molecule trajectories and for reading the manuscript.

This work was partially funded by the Center for Advanced Systems Understanding (CASUS), which is financed by Germany’s Federal Ministry of Research, Technology and Space (BMFTR) and by the Saxon Ministry for Science, Culture, and Tourism (SMWK) with tax funds based on the budget approved by the Saxon State Parliament. A.Y. is supported by the Helmholtz Association Initiative and Networking Fund in the frame of Helmholtz AI as well as by the Helmholtz Foundation Model Initiative within the project “PROFOUND”. The authors gratefully acknowledge the computing time provided on the high-performance computer HoreKa by the National High-Performance Computing Center at KIT (NHR@KIT). This center is jointly supported by the Federal Ministry of Education and Research and the Ministry of Science, Research and the Arts of Baden-Württemberg, as part of the National High-Performance Computing (NHR) joint funding program (). HoreKa is partly funded by the German Research Foundation (DFG). D.S.R, K.K.R., B.A, and N.D.P were funded by a core grant of the MRC-Laboratory of Medical Sciences (UKRI MC-A658-5TY10) and a Wellcome Trust Collaborative Grant (206292/Z/17/Z).

## Data availability

The associated data will be made available via a Zenodo repository at the time of publication.

## Code availability

The code for DeepDiffusion and Kymosketch will be made available via GitHub repositories at the time of publication.

## Author Contributions

K.K.R., B.A., and G.D.M. contributed equally to this work. K.K.R, B.A., G.D.M., A.Y., and D.S.R. conceived the study. G.D.M. designed the neural network model. K.K.R., B.A., G.D.M., and N.D.P. tested and benchmarked the model. K.K.R. and B.A. conducted the optical tweezers experiments and analysed the data. S.B. provided computational infrastructure and supervised high-performance computing experiments. K.K.R, B.A., G.D.M., A.Y., and D.S.R. wrote the initial manuscript draft. All authors approved the final draft.

## Materials and Methods

### Formal problem setting

We model the observed localisations for a molecule undergoing facilitated 1D diffusion as an ordered sequence of *L* observations 𝒟 = *{y*_1_, *y*_2_, …, *y*_*L*_*}* sampled from a real-valued empirical process *Y*_*t*_ : ℝ_+_ *→* ℝ at regularly spaced time points 0 *≤ τ*_1_ *< …< τ*_*L*_ *≤ T*, over a fixed time horizon *T* . We assume these observations are conditionally generated from a latent process *X*_*t*_ : ℝ_+_ *→* ℝ governed by the stochastic differential equation

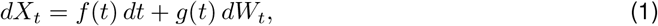

where *f* (*t*) denotes the drift function, and *g*(*t*) denotes the diffusion function, and *W*_*t*_ denotes a standard Wiener process. In our setting, the observed samples are given by *y*_*t*_ = *x*_*t*_ + *ϵ*, where *ϵ* ∼ 𝒩 (0, *σ*^2^) represents additive Gaussian measurement noise, accounting for *LE*.

Our objective is to learn neural estimators *f*_*θ*_(*t* | 𝒟) and *g*_*θ*_(*t* | 𝒟), parameterized by *θ*, that approximate the true drift *f* (*t*) and diffusion *g*(*t*) functions. These estimators are conditioned on the observations 𝒟 and are expected to faithfully reconstruct the dynamics of process *X*_*t*_ on a suitable domain 𝒳 *⊂* ℝ.

Building on the methodologies proposed in prior works [42, 43], we adapt their frameworks to suit the specific structure of our problem. While these prior approaches focus on state-based formulations of stochastic processes, our setting considers a purely time-driven SDE. To accommodate this distinction, we introduce architectural modifications that embed inductive biases aligned with the temporal nature of our dynamics. Additionally, we designed a simplified data generation process that enabled our model to train stably.

### Training Data Generation

In this subsection, we describe the procedure used to generate synthetic datasets of corrupted SDE trajectories, parameterized by smooth, bounded drift and diffusion functions. Building on the general generative model outlined in Eq. 1, we propose using truncated Fourier series with randomized coefficients. Specifically, the drift and diffusion functions are sampled as follows:

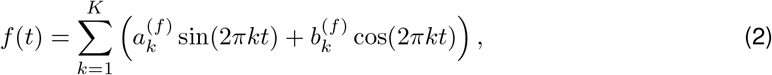

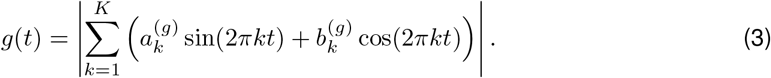

The coefficients 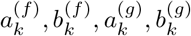 are sampled uniformly from a bounded interval [*−α, α*], where *α* is chosen to ensure that the resulting functions remain within a stable amplitude range. This boundedness not only improves numerical stability during SDE simulation but also promotes regularity in the learned dynamics.

For the diffusion function *g*(*t*), we further enforce positivity by applying an element-wise absolute value, ensuring that the diffusion term in the SDE remains well-defined. This modeling choice aligns with practical scenarios where diffusion is strictly non-negative and varies smoothly with time. The resulting functions are expressive yet well-behaved, making them suitable for simulating a wide range of stochastic systems under controlled scenarios.

To further improve the dataset, we additionally add stepwise training functions for the drift and diffusion coefficients, which are generated by representing each coefficient as a sum of randomly placed rectangular pulses. For every synthetic sample, the function is initialized on a fixed temporal grid, then a random number of rectangles is drawn with random start locations, widths, and amplitudes. Each rectangle contributes a constant value over its support, and overlapping rectangles add together, producing piecewise-constant profiles with abrupt jumps and plateaus. Separate rectangle-generated profiles are used for the drift and diffusion coefficients, giving the training set a broad range of discontinuous or sharply varying dynamics before stochastic trajectories are simulated from them.

To simulate trajectories of the latent stochastic process, we assume a fixed initial condition *x*_0_ = 0 for all generated paths. Given the drift *f* (*t*) and diffusion *g*(*t*) functions sampled as described above, we numerically integrate the stochastic differential equation using the Euler–Maruyama method with a discretization step size ∆*t*. This yields discrete-time samples of the latent process 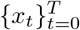 over the time horizon [0, *T* ]. The Euler–Maruyama update rule is given by

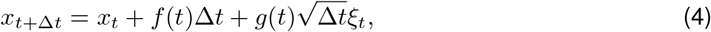

where *ξ*_*t*_ ∼ 𝒩 (0, 1) is standard Gaussian noise. Experimental data is typically noisy due to localisation errors. To account for this, we add an additive Gaussian noise to the simulated path, given by *y*_*t*_ = *x*_*t*_ + *ϵ*, where *ϵ* ∼ 𝒩 (0, *σ*^2^). The noise variance *σ*^2^ may be tuned relative to the scale of the latent dynamics to simulate varying levels of signal-to-noise ratios.

### Model Architecture and Parameterization

Inspired by the design principles introduced in [43], we propose an efficient architecture to process long input sequences arising from fine-grained temporal discretizations over a large temporal horizon. Since the latent SDE is purely time-driven, we utilize a one-dimensional convolutional neural network with residual connections and temporal pooling to compress the observed paths into a compact representation (Figure 2a). This design allows the model to remain tractable even when the number of time steps is large. To ensure robustness across varying discretization granularities, we feed both the observed realizations and their corresponding time grid as inputs. These are processed jointly to yield a sequence of time-dependent latent features.

The compressed output is then passed through a unidirectional LSTM, and we extract the final hidden state **h**_*T*_ as a compact summary of the trajectory. We refer to this hidden state as the path embedding [*X*], which serves as the conditioning context for estimating the drift and diffusion functions.

Formally, given *n* realizations 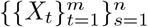 sampled at discrete time points 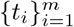, we compute the embedding as:

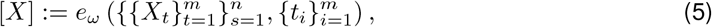

where *e*_*ω*_ denotes the encoder network parameterized by *ω*. Given a query point *t*, the drift and diffusion functions are then evaluated as:

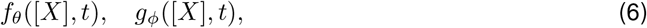

where *θ* and *ϕ* denote the neural parameters of the drift and diffusion networks, respectively. This conditional formulation enables the model to adapt to different observed dynamics while maintaining generalization across paths and time resolutions.

### Optimization Objective

To train the model, we minimize a divergence between the estimated drift and diffusion functions (*f*_*θ*_, *g*_*ϕ*_) and the true functions (*f, g*) over a predefined domain 𝒳 ⊂ ℝ with time horizon *T* . Following standard practice in the neural operator literature [44], we adopt a mean-squared error criterion as our primary divergence measure. The pointwise loss is defined as:

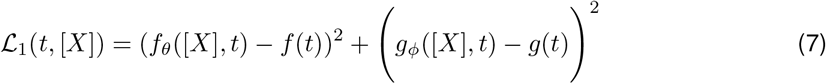

This divergence is evaluated by sampling *t* ∼ 𝒰 (*T* ), where 𝒰 denotes the uniform distribution over the domain.

Several works have shown that reweighting the divergence introduces stability in training and improves the estimated likelihood [42, 44, 45]. To account for local contributions that dominate the integral. Therefore following [42] we introduce an auxiliary learnable function *U*_*v*_(*t*, 𝒟), implemented as a third neural head. The final training objective combines the divergence with an uncertainty-aware reweighting:

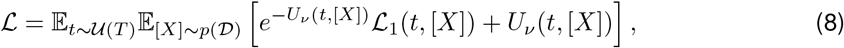

where *p*(𝒟) is the probability distribution over the simulated dataset previously described.

### Testing Synthetic Data

To test the accuracy and precision of DeepDiffusion, synthetic datasets, where the ground truth is already knowm, were generated. Each dataset consisted of 100 displacement vs time trajectories, with the true *D* spanning the experimentally accessible range of 10^*−*4^ to 1 µm^2^/s. The interval time between each data-point was 0.1 s. For each dataset, a localisation error (*LE*) term was further added as Gaussian noise, with widths ranging from 0–300 nm.

For the homogeneous dataset, each trajectory was 30 s long and consisted of a single constant *D*, ranging from 10^*−*4^ to 1 µm^2^/s.

For the heterogeneous case, four distinct datasets were used. The first two consisted of 30 s long trajectories, with single change in *D* at the mid-point of the trajectory, either from a high initial *D*_*fast*_ to a lower final *D*_*slow*_, or from a lower initial *D*_*slow*_ to a high final *D*_*fast*_. The other two datasets consisted of 60 s long trajectories, with two transitions after 20 s and 40 s. The transitions were either from a high *D*_*fast*_ to a lower *D*_*slow*_ and back, or from a lower *D*_*slow*_ to a high *D*_*fast*_ and back. In all four cases, the high *D*_*fast*_ was set to 1 µm^2^/s, and the lower *D*_*slow*_ ranged from 3*×*10^*−*3^ to 0.3 µm^2^/s.

DeepDiffusion was run on each dataset to estimate the corresponding instantaneous *D*s. The relative error (in percentage) was calculated using

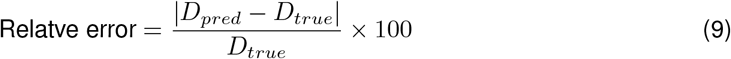

where *D*_*pred*_ is the *D* predicted by DeepDiffusion and *D*_*true*_ is the ground-truth *D*.

### Rolling Diffusion Benchmark

To compare the DeepDiffusion model with contemporary techniques, the rolling diffusion algorithm, as previously described [12], was used as a benchmark. Briefly, MSD *vs* lag-time fits were computed on rolling windows of the data, using a window size of 50 frames, and a linear fit was computed over time delays (tau) of 1 through 5 frames (Figure 1). This window size was optimised to return the least error in terms of precision and blurring over transitions. The diffusion coefficient for each window was subsequently computed from the gradient of the linear fit with the following equation:

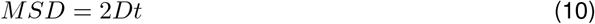

At high *LE*s and low *D*s, the gradient of the MSD against lag-time fit can sometimes be negative. As *D*s cannot be negative (and negative values are especially troublesome to represent on a log scale), these negative values were excised from the data for further analysis. However, for the purpose of display, negative *D*s were replaced with values linearly interpreted from neighbouring points. The relative error for the rolling-diffusion method was calculated in the same manner as for DeepDiffusion above.

### Preparation of DNA

The DNA substrates used in the optical tweezers experiments were prepared as previously described [12, 46].

Briefly, for dsDNA construct, biotin was added to both ends of bacteriophage *λ* genomic DNA (Thermo Scientific) using Klenow polymerase exo^*−*^ (New England Biolabs). The polymerase reaction was carried out by incubating 4 nM of *λ* DNA with 100 *µ*M dGTP, 100 *µ*M dTTP, 80 *µ*M biotin-14-dATP, 80 *µ*M biotin-14-dCTP, and 5 U of polymerase in NEB2 buffer at 37°C for 30 min, then 75°C for 15 min, and subsequently cooling on ice.

For the gapped ssDNA-dsDNA construct, *λ* DNA was end-capped with two biotinylated ssDNA oligos using T4 ligase (New England Biolabs) to form dsDNA with closed biotinylated ends [47]. This product was subsequentely site-specifically nicked at two locations in the same strand using CRISPR-Cas9_D10A_ nickase (IDT). The nicked product was incubated with proteinase K (New England Biolabs) at 56°C for 15 min to remove any nickase bound to the ssDNA-dsDNA junction. The ssDNA gap was formed *in situ* during the experiments using force-induced melting, as described previously [46].

### CTFM experiments

Single-molecule CTFM experiments were performed on a C-trap (Lumicks), combining optical tweezers, confocal fluorescence micrscopy, and microfluidics. The five-channel microfluidic flowcell was passivated using 0.5% (w/v) Pluronic F-127 in PBS and, subsequently for channels 3, 4, and 5, with 1 mg/ml BSA.

D2-I data were collected for a previous study of D2-I diffusion on DNA [12]. These experiments were carried out as previously described. Briefly, the following were introduced into the respective channels by flow: i) 0.005% (w/v) streptavidin-coated polystyrene beads (∼4.4 *µ*m, Spherotech) in channel 1; ii) ∼2pM of biotinylated DNA construct in channel 2; iii) D2-I Experimental Buffer (20 mM HEPES pH=7.5, 75 mM NaCl, 0.5 mg/ml BSA, 1 mM TCEP) in channel 3; iv) 5 nM fluorescently labelled D2-I (with LD555 and LD655, Lumidyne Technologies, on D2 and I, respectively) in D2-I Experimental Buffer in channel 4; v) 800 pM eGFP-RPA in D2-I Experimental Buffer in channel 5. The streptavidin-coated beads were trapped optically in channel 1 (with trap stiffness of 0.2–0.3 pN/nm) and moved to channel 2 to tether DNA, which was subsequently verified in channel 3. The ssDNA gap was generated at this stage. The DNA tether was briefly incubated in the protein channels, before being imaged in channel 3 using confocal lasers (488 nm, 532 nm, and 638 nm). Kymographs were acquired using a pixel dwell-time of 0.1 ms per pixel.

INO80 data were collected using core human INO80 (purified as described previously [48]) with an Atto647N label via a ybbR tag [49] on Ies6. The following were introduced into the respective channels by flow: i) 0.005% (w/v) streptavidin-coated polystyrene beads (∼4.4 *µ*m, Spherotech) in channel 1; ii) ∼2pM of biotinylated lambda DNA in channel 2; iii) INO80 Buffer 1 (50 mM Tris pH=7.8, 100 mM KCl, 2 mM MgCl_2_, 1 mM EDTA, 0.5 mg/ml BSA) in channel 3; iv) 1 nM fluorescently labelled hINO80 (as above) in INO80 buffer 1. The streptavidin-coated beads were trapped optically in channel 1 (with trap stiffness of 0.2–0.3 pN/nm) and moved to channel 2 to tether DNA, which was subsequently verified in channel 3. The DNA tether was briefly incubated in channel 4, before being imaged in channel 3 using confocal lasers (638 nm). Kymographs were acquired using a pixel dwell-time of 0.1 ms per pixel.

### Experimental data analysis

Single-molecule displacement *vs*. time trajectories were extracted from kymographs using KymoSketch (see Supplementary Information). Manually identified and traced molecules were sub-pixel localised using a 7 pixel Gaussian refinement window. The extracted localisations were subsequently used to generate displacement *vs*. time trajectories which were further analysed.

Trace-averaged diffusion coefficients were estimated by fitting the corresponding MSD *vs*. lag-time curves with a line and using Eq. 10. The rolling window estimate of instantaneous *D* was similarly calculated using linear fits of MSD *vs*. lag-time curves calculated for rolling windows of 50 frames. For DeepDiffusion, the space and time symmetry of the Brownian diffusion process was utilised to inform on the uncertainty within the model. The model was run on the trajectory four times, in the unaltered ‘forward’ direction (the ‘forward’ model), with the time-axis reversed (the ‘reverse-time’ model), with the space-axis reversed (the ‘reverse-space’ model), and with both axes reversed (the ‘reverse-space-time’ model). The mean of the four iterations was taken as the estimated *D*, while the standard deviation of the iterations reported on the model uncertainty.

For the INO80 diffusion experiments, *D* distributions were modelled with Gaussian mixture models (GMM) [50], using a maximum-likelihood approach, as previously described [51].

## Supplementary Figures

**Supplementary Fig. S1.**
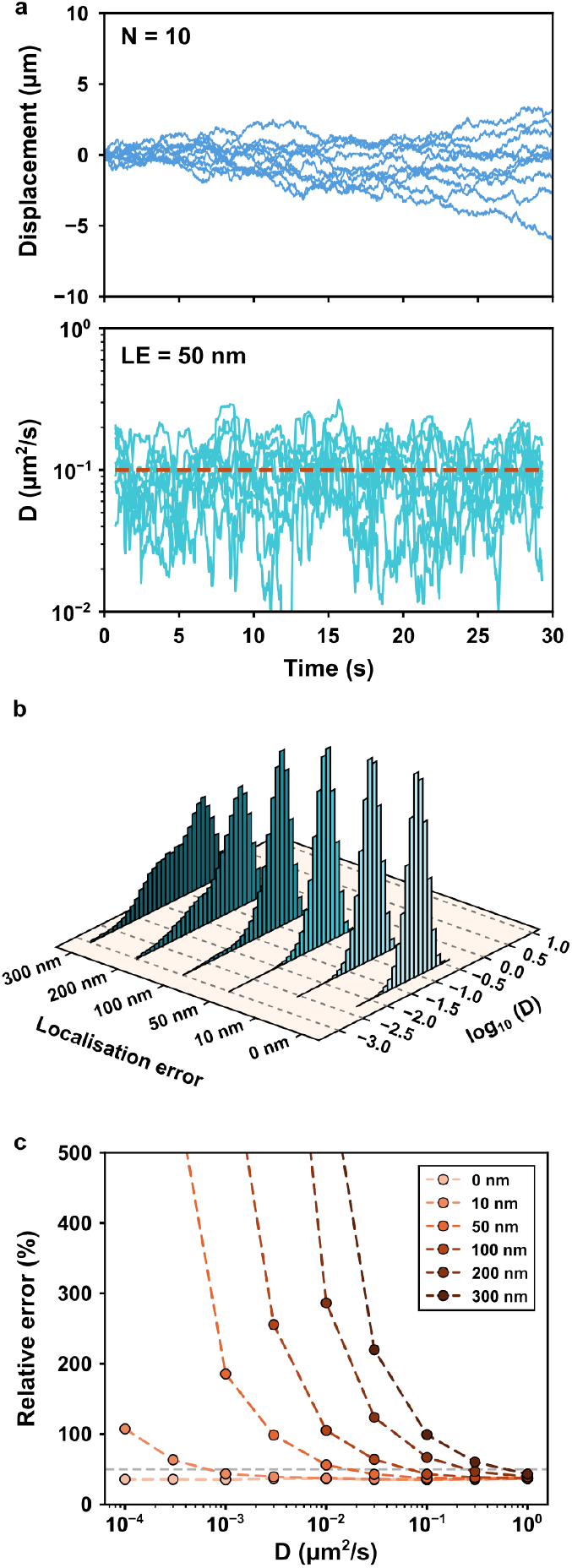
Benchmarking rolling-window analysis using synthetic homogeneous datasets. **a**, Representative single-molecule trajectories (blue) with constant *D* (*top*), along with estimated (cyan) and true (red) instantaneous *D* (*bottom*). **b**, Estimated *D* histogramss with increasing *LE* for *D* = 0.1 µm^2^/s. **c**, Relative errors of estimated *D* plotted for varying constant *D*s and *LE*.

**Supplementary Fig. S2.**
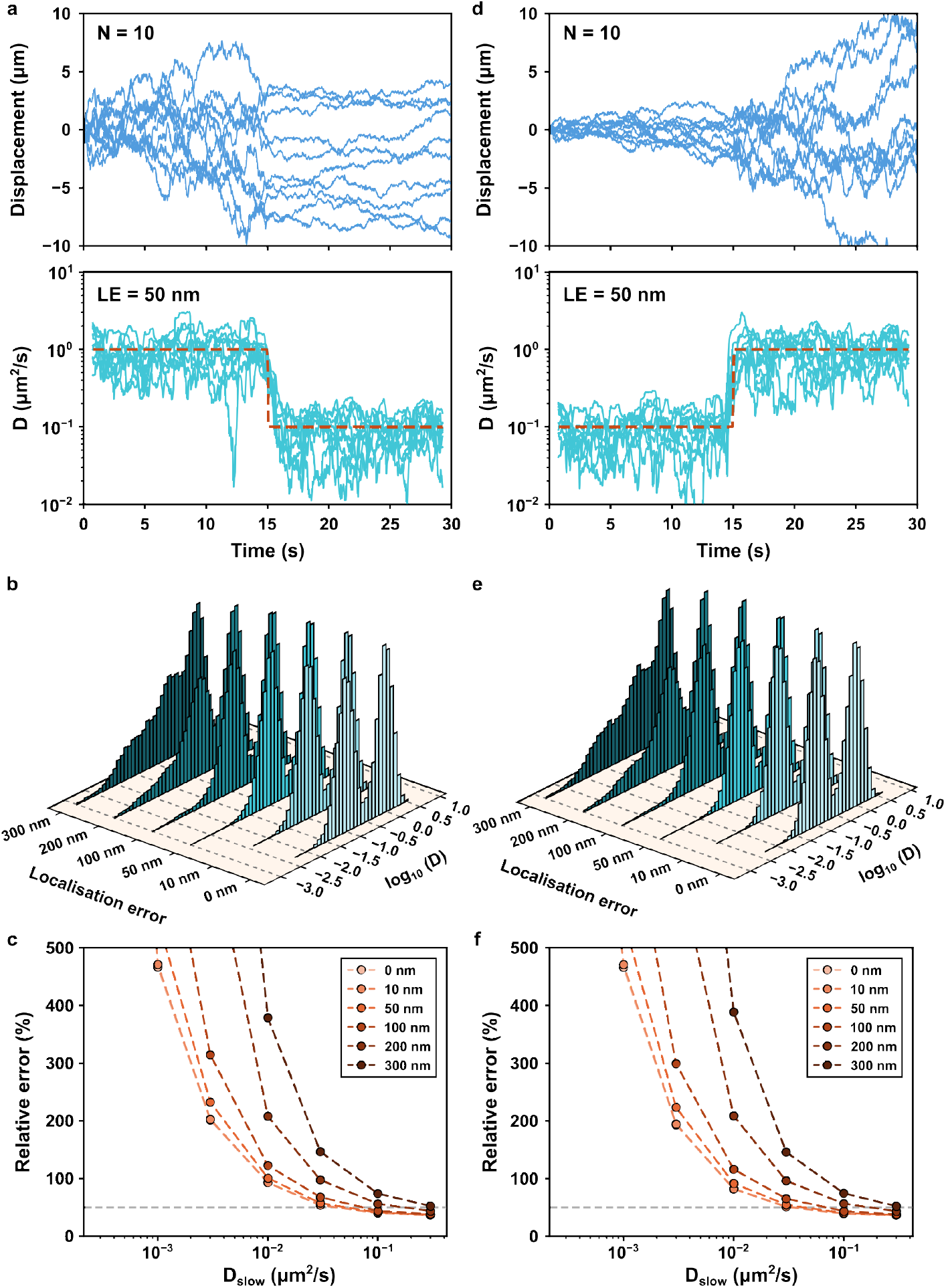
Benchmarking rolling-window analysis using synthetic heterogeneous datasets. **a**, Representative single-molecule trajectories (blue) with a single decreasing change in *D* (*top*), along with estimated (cyan) and true (red) instantaneous *D* (*bottom*). **b**, Estimated *D* histograms with increasing *LE* for initial *D*_*fast*_ = 1 µm^2^/s and final *D*_*slow*_ = 0.1 µm^2^/s. **c**, Relative error of estimated *D*s plotted for varying final *D*_*slow*_s and *LE*. **d**, Representative single-molecule trajectories (blue) with a single increasing change in *D* (*top*), along with their estimated (cyan) and true (red) instantaneous *D* (*bottom*) **e**, Histograms of estimated *D*s with increasing *LE* for datasets with an initial *D*_*slow*_ = 0.11 µm^2^/s and a final *D*_*fast*_ = 1 µm^2^/s. **f**, Plot of the relative errors of estimated *D*s for varying initial *D*_*slow*_s and *LE*.

**Supplementary Fig. S3.**
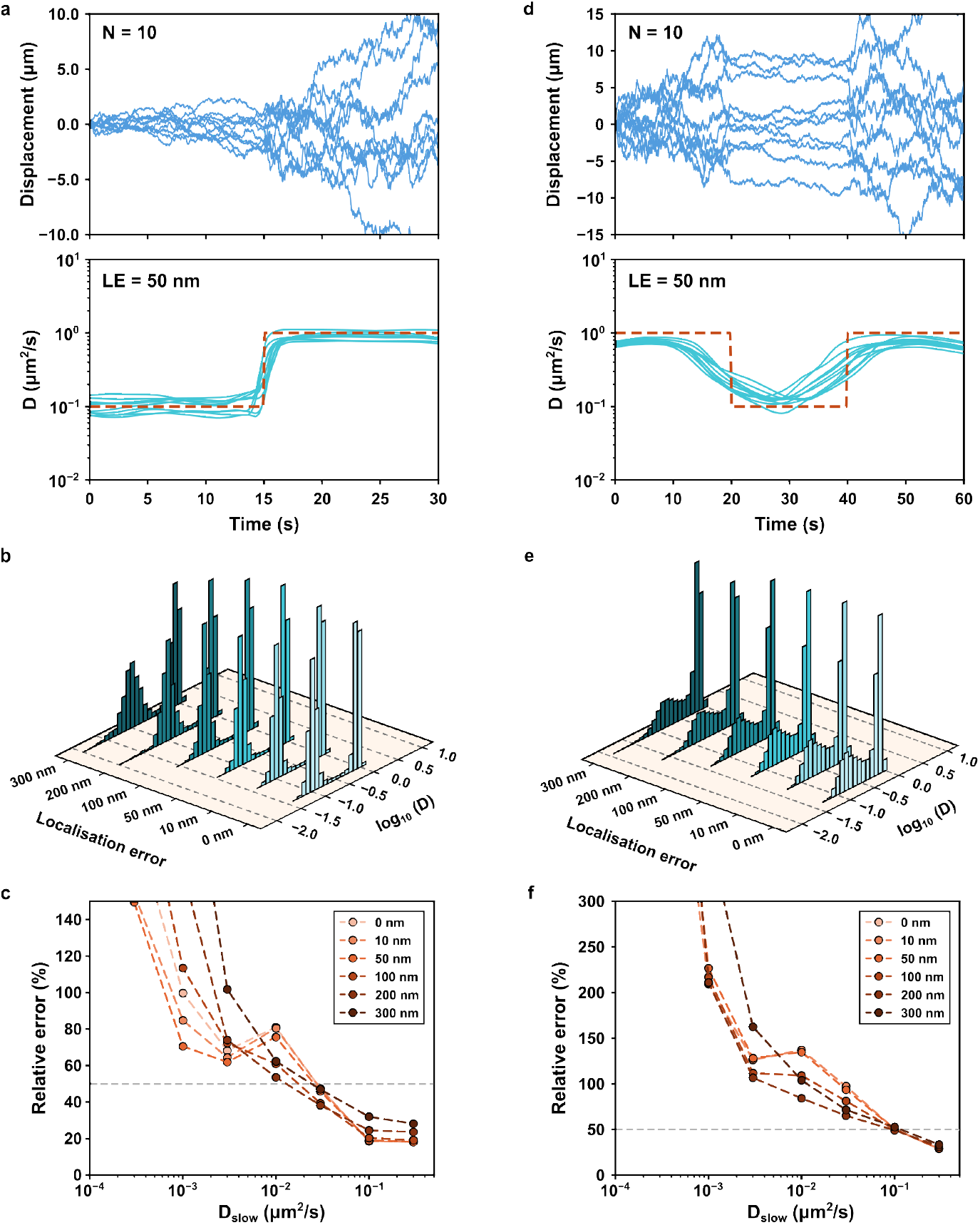
Evaluation of DeepDiffusion using synthetic heterogeneous datasets. **a**, Representative single-molecule trajectories (blue) with a single increasing change in *D* (*top*), along with their predicted (cyan) and true (red) instantaneous *D* (*bottom*). **b**, Predicted *D* histograms with increasing *LE* for initial *D*_*slow*_ = 0.1 µm^2^/s and a final *D*_*fast*_ = 1 µm^2^/s. **c**, Relative errors of predicted *D*s plotted for varying initial *D*_*slow*_s and *LE*. **d**, Representative single-molecule trajectories (blue) with two changes in *D* (*top*), along with their predicted (cyan) and true (red) instantaneous *D* (*bottom*). **e**, Predicted *D* histograms with increasing *LE* for an initial and final *D*_*fast*_ = 1 µm^2^/s and an intermediate *D*_*slow*_ = 0.1 µm^2^/s. **f**, Plot of the relative errors of predicted *D*s for varying intermediate *D*_*slow*_s and *LE*.

**Supplementary Fig. S4.**
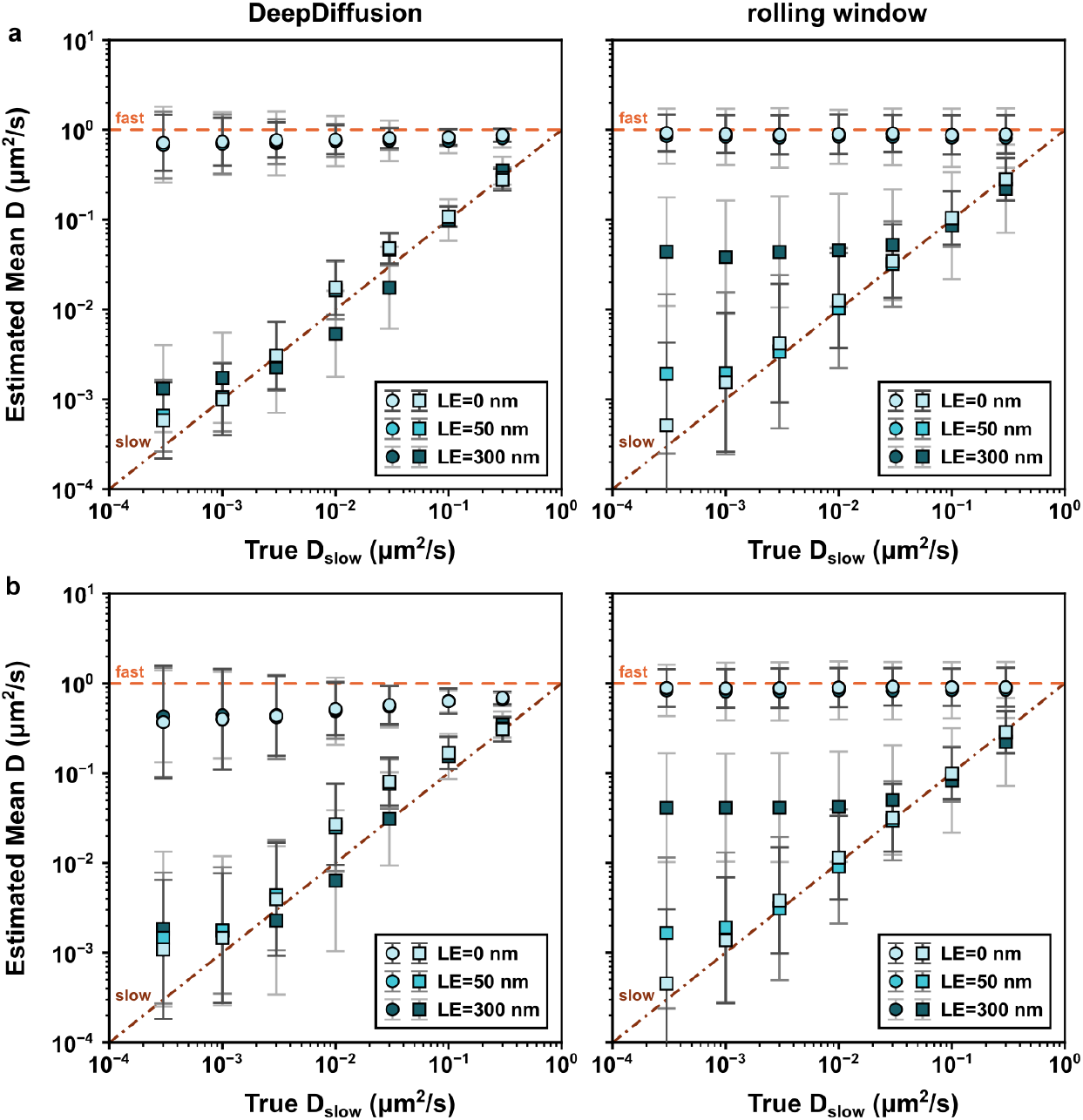
Evaluation of DeepDiffusion bias using synthetic heterogeneous datasets. **a**, Mean *D*s (circles and squares for *D*_*fast*_ and *D*_*slow*_, respectively) calculated using DeepDiffusion (left) and rolling window (right) for a single-step decrease from a higher *D*_*fast*_ to a lower *D*_*slow*_, plotted as a function of the true *D*_*slow*_ for different *LE*s. The dashed lines *y* = 1 and *y* = *x* represent the cases of no overall bias for *D*_*fast*_ and *D*_*slow*_ respectively. **b**, ean *D*s (circles and squares for *D*_*fast*_ and *D*_*slow*_, respectively) calculated using DeepDiffusion (left) and rolling window (right) for a two-step change from a higher *D*_*fast*_ to a lower *D*_*slow*_ and back, plotted as a function of the true *D*_*slow*_ for different *LE*s. The dashed lines *y* = 1 and *y* = *x* represent the cases of no overall bias for *D*_*fast*_ and *D*_*slow*_ respectively. Error bars correspond to standard deviations of the *D* distributions.

**Supplementary Fig. S5.**
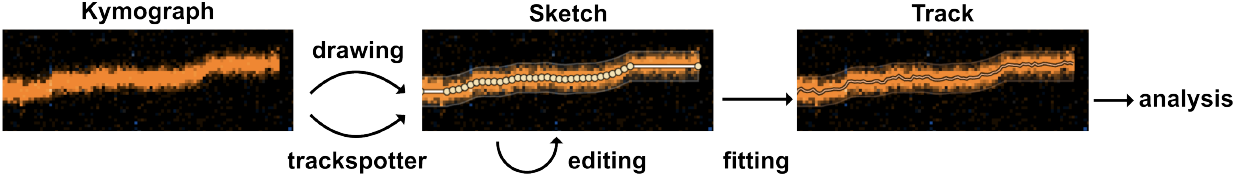
Schematic of the Kymosketch workflow. Kymograph images are loaded into Kymosketch, and preliminary tracks (referred to here as sketches) are produced either via an automated trackspotter algorithm or drawn by the user. These sketches can be editted, using an end finder algorithm to refine the end of the track using photon data, merging tracks the trackspotter misidentified as separate or splitting tracks that the track spotter miscombined. Fitting algorithms (centroid and MLE) can then be used to refine the sketch into a finalised track for further analysis, either with conventional methods or deep-diffusion

## Supplementary Information

### Kymosketch Algorithms

#### Automated Track Detection

Track detection proceeds in two phases, per-column peak detection followed by greedy trajectory linking.

#### Peak detection

For each image column (*i*.*e*., each time frame) of the kymograph, a three-pixel sum along the spatial axis is used to reduce noise (expect for the edges). After subtracting the profile mean, the signal is smoothed with an 11-point, third-order Savitzky–Golay filter (coefficient array = [*−*36, 9, 44, 69, 84, 89, 84, 69, 44, 9, *−*36]*/*429), which preserves peak shape better than a moving average. Peaks are identified as strict local maxima exceeding a threshold, then refined by greedy non-maximum suppression with a minimum spacing of 3 px.

#### Trajectory linking

A greedy nearest-neighbour linker scans columns left to right. Each seed peak is refined via an intensity-weighted centroid within a local window (background-subtracted using the local minimum). Peaks in successive columns are linked if the displacement is less than a step parameter; otherwise the trajectory terminates. Final trajectories are trimmed by 2 points at each end and discarded if shorter than a minimum length.

#### Track Endpoint Detection

To locate where a track begins or ends beyond a user-supplied sketch endpoint, we fit an optimal step function (changepoint model) to the intensity profile. A scan range is constructed from 5 columns inside the sketch (baseline) plus a configurable number of columns beyond the endpoint. At each column, the intensity is summed over a vertical window of size 2*w* + 1 centred on the track position.

Using a cumulative-sum formulation for *O*(1) segment-mean evaluation, we exhaustively test every candidate breakpoint *k* (minimum segment length 3) and score it as

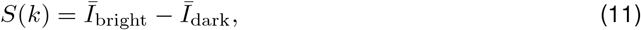

where the bright/dark assignment depends on direction. The breakpoint is accepted only if the intensity drop exceeds 30% of the bright-side mean.

#### Centroid Fitting

Sub-pixel track positions are obtained by iterative intensity-weighted centroid refinement. To handle overlapping tracks, nearby trajectories (separation *≤* 2*w*) are clustered and each track’s fitting region is bounded by the midpoints between neighbours.

#### Iteration 0 (pure centroid)

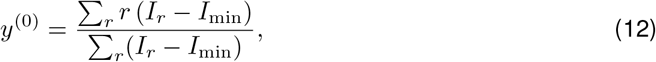

where *I*_min_ is the minimum intensity in the window.

#### Iterations ≥ 1 (Gaussian-weighted centroid)

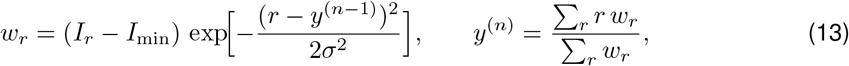

with *σ* = *w/*2. The Gaussian kernel suppresses bias from asymmetric tails and nearby contaminants. Results are clamped to overlap and image bounds after each iteration.

#### Poisson–Gaussian Maximum-Likelihood Fitting

Each column’s intensity profile is modelled as a sum of Gaussians plus a constant background:

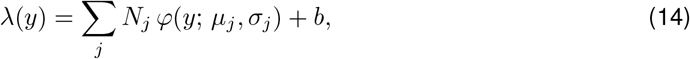

where *φ* is the normal PDF, *N*_*j*_ is the integrated photon count (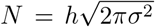 for peak height *h*), *µ*_*j*_ the centre position, *σ*_*j*_ the PSF width, and *b* a shared background. Pixel counts are assumed Poisson-distributed, giving the log-likelihood

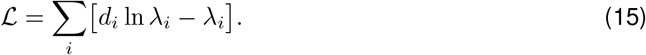

#### Fisher scoring with Levenberg–Marquardt damping

Parameters are optimised iteratively (up to 30 steps). The Jacobian entries are

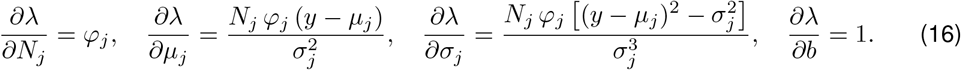

The score vector and expected Fisher information are

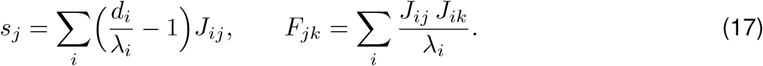

LM damping is applied as *F*_*jj*_ += 0.01 (*F*_*jj*_ + 10^*−*4^), and the system *F δ* = *s* is solved via Gaussian elimination with partial pivoting. A backtracking line search (halving up to 20 times) ensures monotonic likelihood increase; iteration stops when *∥s∥*_2_ < 10^*−*6^.

Parameter bounds enforce *N* > 0, *σ* ∈ [0.5, 10] px, and restrict *µ* to the inner half of the fitting window so that data always flank the peak on both sides.

